# rDNA nascent transcripts promote a unique spatial organization during mouse early development

**DOI:** 10.1101/2021.04.20.440564

**Authors:** Martine Chebrout, Maimouna Coura Kone, Habib U. Jan, Marie Cournut, Martine Letheule, Renaud Fleurot, Tiphaine Aguirre-Lavin, Nathalie Peynot, Alice Jouneau, Nathalie Beaujean, Amélie Bonnet-Garnier

## Abstract

During the first cell cycles of the early development, the chromatin of the embryo is highly reprogrammed alongside that embryonic genome starts its own transcription. The spatial organization of the genome is a major process that contributes to regulating gene transcription in time and space, however, it is poorly studied in the context of early embryos. To study the cause and effect link between transcription and spatial organization in embryos, we focused on the ribosomal genes, that are first silent and begin to transcribe during the 2-cell stage in mouse. We demonstrated that ribosomal sequences are spatially organized in a very peculiar manner from the 2-cell to the 16-cell stage with transcription and processing of ribosomal RNAs excluding mutually. Using drugs inhibiting the RNA polymerase I, we show that this organization, totally different from somatic cells, depends on an active transcription of ribosomal genes and induces a unique chromatin environment that favors major satellite sequences transcription after the 4-cell stage.

## Introduction

In eukaryotes, the spatial organization of the genome within the interphasic nuclei is not random. DNA fluorescent in situ hybridization (DNA-FISH) experiments have demonstrated that chromosomes occupy a specific nuclear position called territories ((Cremer and Cremer, 2001) and that transcriptional activity may dictated gene positioning reviewed in (Meaburn, 2016). The three-dimension (3D) organization of chromatin acts as a key component of the cell identity and can be correlated with highly differentiated cell types (Solovei et al., 2009). However, the relationships between genome organization and gene transcription is still a matter of debate, albeit after extensive studies reviewed in (van Steensel and Furlong, 2019). For a decade, chromosome conformation capture techniques (3C, 4C and Hi-C) offer the possibility to study whole-genome organization, long-distance interaction, and loops between genomic regions or loci (reviewed in (Bonev and Cavalli, 2016). But depending on the biological model, DNA-FISH and microscopy are still relevant and at least complementary to these approaches (Szabo et al., 2020) allowing single-cell analysis in intact nuclei (Gelali et al., 2019) and localization of repeated sequences such as pericentromeric and centromeric regions of ribosomal DNA (rDNA) genes (Maiser et al., 2020; Mayer et al., 2005).

As for mammalian embryos, 3D-FISH of pericentromeric regions (comprising minor and major satellite sequences, (Guenatri et al., 2004)) has been extensively used to demonstrate that these regions undergo large-scale reorganization during the early embryonic development period (reviewed in (Jansz and Torres-Padilla, 2019). We and others have shown that dramatic changes occur throughout the two first cell cycles of mouse preimplantation development concomitantly with the onset of embryonic transcription (also called embryonic genome activation (EGA)) (Aguirre-Lavin et al., 2012; Bonnet-Garnier et al., 2018; Probst et al., 2007). This spatial reorganization of heterochromatin was shown to be required for further development (Casanova et al., 2013; Probst et al., 2010), highlighting its importance. Remarkably, in 1-cell and early 2-cell mouse embryos, major (and minor) satellite sequences surround dense, spherical structures called nucleolar precursor bodies (NPBs -(Fléchon and Kopecny, 1998)). While, the NPBs were first described as a seed for embryonic nucleolus to settle (Zatsepina et al., 2003), it is now believed that they serve rather as a structural platform anchoring heterochromatin and allowing its remodeling (Fulka and Langerova, 2019). At the time of EGA, satellite sequences indeed progressively disconnect from these NPBs while forming round shape clusters as found in somatic cells and called chromocenters (Aguirre-Lavin et al., 2012). Recently, (Hamdane et al., 2016) have shown that embryos lacking the nucleolar protein UBF also lack NPBs and display an abnormal distribution of heterochromatin. However, the precise relationship between these sequences and the NPBs remains unclear.

The inner organization of NPBs has been thoroughly investigated (Baran et al., 1995; Baran et al., 2001; Koné et al., 2016) by immunofluorescent staining of nucleolar proteins such as Upstream Binding Transcription Factor (UBTF), Fibrilarin, B23/Nucleophosmin 1 (NPM1), and Nopp140/NOLC1 (Nucleolar and Coiled-body phosphoprotein 1) showing dynamic redistribution of the different nucleolar compartments (mainly the dense fibrillar component, (DFC) and the fibrillar center, (FC)) between the 2-cell and blastocyst stages (time of implantation). The NPBs also structurally support the ribosomal genes (rDNA) (Romanova, 2006). Remarkably, reinitiating of ribosomal transcription (rRNA synthesis) has been shown - by BrUTP incorporation (Zatsepina et al., 2003) - to occur in NPBs at the end of the 2-cell stage, when pericentromeric repeats undergo massive reorganization. It could be that the chromatin state of both rDNA and pericentromeric sequences influence each other as described in Embryonic stem cells (Guetg et al., 2010; Savić et al., 2014). Little is known regarding the spatial organization and expression of rDNA genes in the nuclei of mouse embryos.

In the somatic nuclei, the ribosomal genes are transcribed in the nucleolus by the RNA polymerase I (RNA pol I) in a long precursor transcript called 47S pre-rRNA, that will be cleaved by several exonucleases to separate the internal transcribed spacers 1 (ITS1) and 2 (ITS2) and the 5′ and 3′ external transcribed spacers (5′-ETS and 3′-ETS) from the mature ribosomal RNAs (rRNAs): 18S, 5.8S, and 28S (Henras et al., 2015). These rRNAs are associated with ribosomal proteins in pre-ribosomes particles, processed/matured by several proteins, and exported in the cytoplasm to form the small and large ribosome subunits (Mullineux and Lafontaine, 2012). Two proteins: the UBF and the Selective factor 1 (SL1) are required to load the Pol I complex and initiate transcription of the rDNA (Moss et al., 2019). Experimental disruption of the rDNA transcription can be done using either Actinomycin D that intercalates into DNA at the actively transcribed rDNA sites and inhibits pre-rRNA chain elongation (Schöfer et al., 1996) or CX-5461 that specifically inhibits Pol I transcription of rDNA genes by selectively targeting the SL1 transcription factor (Drygin et al., 2011). Studies have shown that these inhibitors are able to affect nucleolar structure organization,reviewed in (Grummt, 2013; Mangan et al., 2017; Potapova and Gerton, 2019).

With the use of these specific inhibitors, we will address the organization of rDNA in time and space with regards to major satellite sequences during early embryonic development by 3D-FISH. We will also analyze expression of the various rRNA transcripts to decipher the causal relationship between the spatial organization and the expression changes of these sequences.

## Results

### Ribosomal genes (rDNA) 3D organization is linked to their transcription status during preimplantation development

To determine the spatial distribution of rDNA repeats concomitantly with major satellite sequences in mouse early embryo nuclei, 3D-DNA-FISH was performed using probes specific to mouse rDNA repeats (Akhmanova et al., 2000; van de Nobelen et al., 2010) and major satellite sequences (Aguirre-Lavin et al., 2012), respectively. Preservation of the 3D structure of the whole embryos allowed us to do in-depth localization of these sequences from the 1-cell to the blastocyst stage (Fig. 1A and 1C, Fig. S1A and S1B).

**Figure 1:**
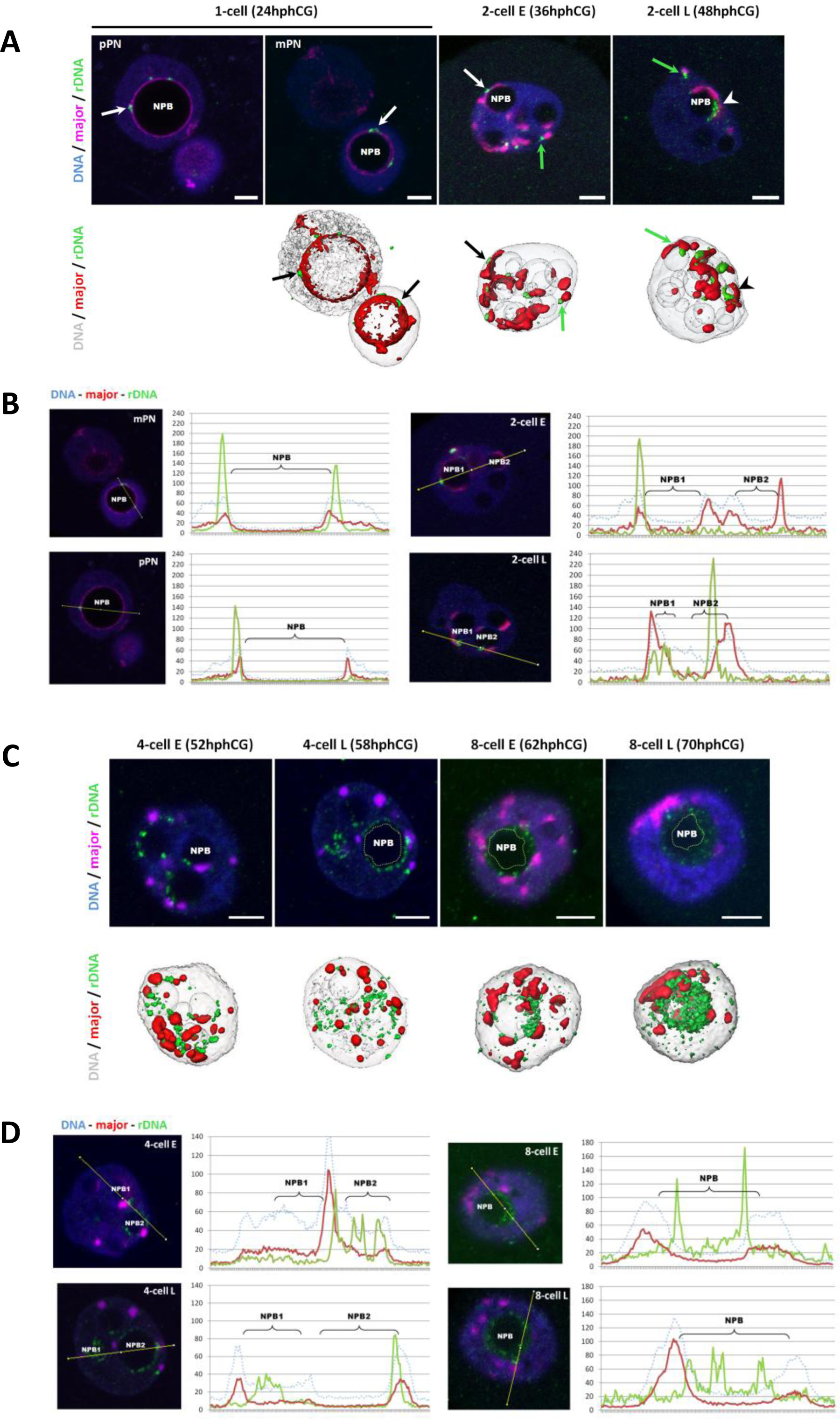
3D organization of ribosomal sequences (rDNA) is correlated with their transcriptional state. **(A)** 3D organization of rDNA and major satellite sequences at 1-cell and 2-cell stages detected by 3D DNA-FISH using specific probes. <u>Upper panel :</u> single z-section of a representative nucleus of 1-cell (24hphCG), early (E, 36hphCG) and late (L, 48hphCG) 2-cell stages. <u>Lower panel:</u> Amira 3D representation of each nucleus. White (or black, lower panel) arrows indicate clustered rDNA FISH signal juxtaposed to major satellite signal at NPBs surface; green arrows indicated rDNA juxtaposed to round shape major satellite signal and white (or black, lower panel) arrowhead indicate pearl necklace rDNA signal at NPBs surface. **(B)** Localization of rDNA and major satellite signals in regards to NPBs boundaries at 1-cell and 2-cell stages. <u>Left panel:</u> single z-section of the nucleus with the line corresponding to the Intensity profile measurement in Fiji. <u>Right panel:</u> Fluorescence intensity plot profile along the line across multiple channels (blue dot line for DAPI, the green line for rDNA signal, and red line for major satellite signal). **(C)** 3D organization of rRNA genes from 4-cell to 8-cell stages detected by 3D DNA-FISH. rDNA FISH signal expands in the inner of the NPBs up to fulfil it completely. <u>Upper panel:</u> z-section of a representative nucleus of early (E) or late (L) 4-cell and 8-cell stage. <u>Lower panel:</u> Amira 3D representation of each nucleus. **(D)** Localization of rDNA and major satellite signals in regards to NPBs boundaries at 4-cell and 8-cell stages. <u>Left panel:</u> single z-section of the nucleus with the line corresponding to the Intensity profile measurement in Fiji. <u>Right panel:</u> Fluorescence intensity plot profile along the line across multiple channels (blue dot line for DAPI, the green line for rDNA signal, and red line for major satellite signal).DNA is in blue or grey, rDNA in green, and major satellite sequences in magenta. *Scale bar = 5µm*. mPN: maternal Pronucleus; pPN : paternal Pronucleus and NPB: Nucleolar Precursor Body; h phCG : hours post-injection of human Chorionic Gonadotrophin, major: mouse major satellite sequences, rDNA: ribosomal DNA.

As expected during the 1-cell stage, major satellite sequences progressively formed a ring that surrounded NPBs (Fig. S1A, (Aguirre-Lavin et al., 2012). rDNA signals correspond to large foci (Fig. 1A and Fig. S1A) and are always associated with major satellite sequences irrespective of the origin of the pronucleus (mPN or pPN). At the 2-cell stage, rDNA FISH signals have a small spots shape close to large signal of major satellite sequences and can be divided into two categories: (i) some of them are not associated with NPBs (green arrows in Fig. 1A) and (ii) some of them are embedded within the ring of major satellite sequences surrounding the NPBs (black and white arrows in Fig. S1A). Interestingly, at the late 1-cell and early 2-cell (2-cell E) stages, these rDNA signals are localized at the outer edge of the NPBs (Fig. 1A) as demonstrated by plot profiles of fluorescence intensities for DAPI, major sequences, and rDNA signals across NPBs (mPN, pPN and 2 -cell E in Fig. 1B). At the end of the 2-cell stage (2-cell L) concomitantly with the embryonic genome activation (Zatsepina et al., 2003), rDNA FISH signals change in shape and localization: forming pearl necklace structures (arrowhead in Fig. 1A) juxtaposed to major satellites and extending inside the NPBs (green pick inside NPB2, plot profile at 2-cell L in Fig. 1B).

From the early 4-cell to the late 8-cell stage, rDNA signals that were mostly located at the surface of NPBs, lose the pearl necklace shape to form a dispersed cloud of smaller dots, being less and less associated with major satellite sequences (Fig. 1C) and more and more inside the NPBs (as shown by plot profiles drawn across NPBs, Fig. 1D). Finally, the 3D rDNA organization changes one last time, between the 16-cell and the Morula stage: the FISH signals fill the NPBs (16-cell E, 75hphCG in Fig. S1B) and acquire a nucleolus-like structure (Morula) like in differentiated cells (Junera et al., 1995), with no differences between inner cell mass (ICM) and trophectoderm cells (TE) at the Blastocyst stage (Fig. S1B).

Quantification of the number of NPBs per stage revealed that it decreased from the late 2-cell to the 16-cell stage (black line in Fig. S1C) and that the remaining NPBs at the 16- cell stage are mostly those associated with the rDNA signal (dark blue bars in Fig. S1C). We then analyzed how rDNA organization evolves and defined four types of NPBs (Fig. 2A, upper panel): T1 corresponding to NPBs with a small number of round-shape rDNA spots; T2 corresponding to NPBs with larger spots distributed like a pearl necklace, T3 corresponding to NPBs surrounded by a thin cloud of rDNA signals and T4 corresponding to NBPS with a larger cloud of rDNA signals. Analysis of 350 nuclei from 4-cell to 16-cell stages embryos (Fig. 2A lower panel) shows that T1 and T2 NPBs are predominant at the 4-cell stage and decrease significantly up to the late 8-cell stage (p-value < 0.005). On the other hand, T3 NPBs are mostly observed in late 4-cell and early 8-cell stages and T4 NPBs number increases significantly from 8-cell to 16-cell stage (p-value < 0.005, Mann-Whitney U test).

**Figure 2:**
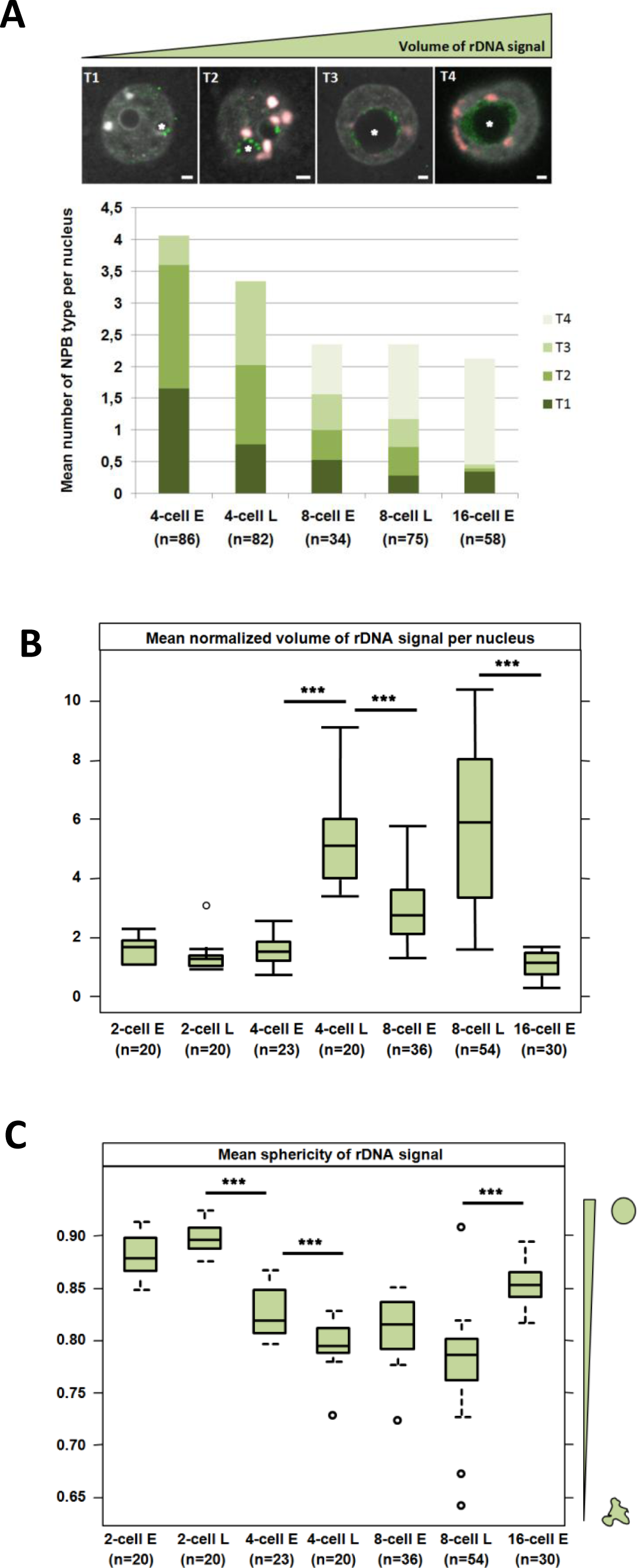
Features of rDNA spatial organization during pre-implantation development. **(A)** Spatial organization rDNA surrounding the NPBs. <u>Upper panel:</u> Four kinds of NPBs can be defined depending on their rDNA pattern: Type 1 (T1) displays small dots of rDNA signal, Type 2 (T2) shows spots distributed like a pearl necklace, Type 3 (T3) harbors a thin cloud of rDNA signal and Type 4 (T4) a large cloud of rDNA signal. Stars indicated the NPBs representative of each type. Scale bar = 5µm. <u>Lower panel:</u> Quantification of the number of each type of rDNA/NPBs pattern per stage at the 4-cell, 8-cell (early (E) and late (L)) and the early 16-cell stage. Mann-Whitney test was used to compare (two by two) the distribution of the type between stages. **, p-value< 0.01; ***, p-value < 0.001. **(B and C)** Quantification of rDNA total volume (B) and average sphericity (C) per nucleus and stage based on the DNA-FISH signal of the ribosomal sequences. All measurements were done using Imaris 9.6 (Oxford Instruments) from the 2-cell to 16-cell stage. Statistical significance was evaluated using a non-parametric multiple comparison test (nparcomp, R package). ***, p-value < 0.001. The number of nuclei examined per stage is indicated below the stage name.

To complete our analysis of rDNA 3D-organization, Imaris software (version 9.6, Oxford Instruments) was used to measure the volume of rDNA FISH signals in whole embryos from the 2-cell to 16-cell stage (Fig. S1D) and the corresponding nuclear volume (stained with DAPI, Fig. S1E). To correct for the potential variation of the nuclear volumes (Fig. S1E), the total rDNA volume in a nucleus was divided by the volume of this nucleus (Fig. 2B). While we did not observe variations at the 2-cell stage, earlier stages have always significantly lower normalized mean volumes than later ones at 4 and 8-cell stages (Fig 2B, S2D) potentially linked with the cell cycle since the chromatin is condensed into chromosomes during mitosis. The late 4-cell, early, and late 8-cell stage displayed larger rDNA volumes than earlier stages, suggesting an important decondensation of the rDNA sequences as visualized on DNA-FISH images (Fig. 1B, lower panel). Remarkably, such decondensation is confirmed by calculating the mean sphericity of rDNA spots (in a given nucleus) at each stage (Fig. 2C). Indeed, 4-cell and 8-cell stage embryos have significantly lower sphericity mean values (from 0.77to 0.82, p-value <0.005) when compared to those of 2-cell and 16-cell stage embryos (0.88 and 0.85), respectively. With regards to these parameters (rDNA volume and sphericity), the 16-cell stage has to be considered apart as the structure of the signal changes dramatically at this stage when compared to previous stages and because nuclei become asynchronous, and early vs. late stage cannot be distinguished anymore.

Altogether, our results show that rRNA sequences change in shape and distribution twice during early development: first between the early and the late 2-cell stage, when rDNA becomes transcriptionally active, and, secondly, between the late 8-cell stage and the 16-cell stage on the commencement of cell differentiation.

### rRNA transcription during early development: a two-step dynamic

To investigate the link between 3D organization of rRNA genes and their transcriptional status, localization of probes specific to the classical rRNA (18S and 28S) and to several pre-rRNA regions that are not retained after complete rRNA processing (5’ETS, ITS1 and ITS2, detailed in Materials and Methods section and Table S1) were studied using RNA-FISH on 3D-preserved embryos (Fig. 3A and 3B).

**Figure 3:**
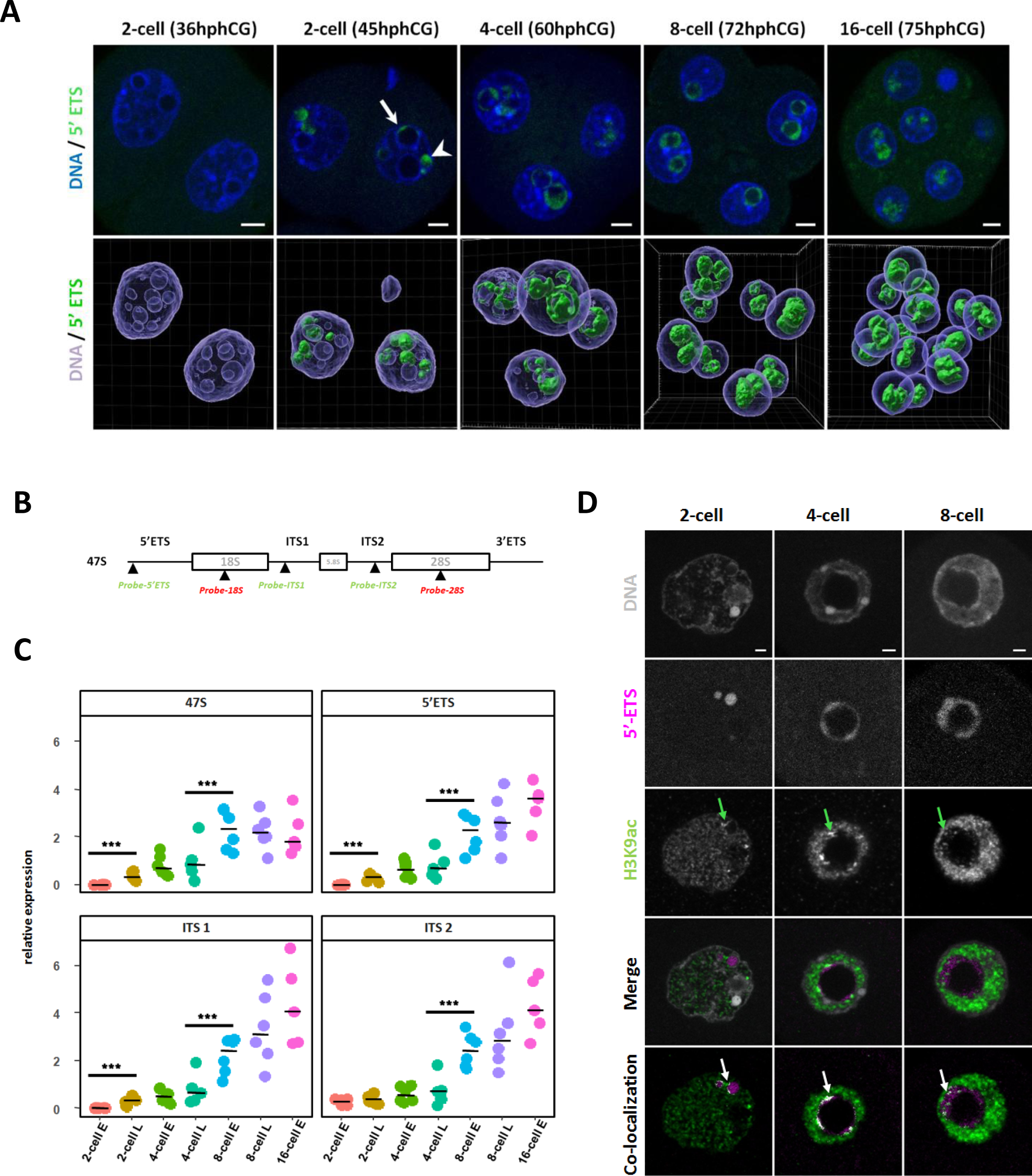
rRNA transcripts organization upon early development. **(A)** rDNA transcripts (green) are visualized by RNA-FISH using a specific probe for the 5’ETS region. <u>Upper panel</u>: representative nuclei of 2-cell (36h and 45hphCG), 4-cell (60hphCG), 8-cell (72hphCG) and 16-cell (75hphCG) stages embryos. Arrow indicates nascent rRNA labeled with a 5’ETS probe at the surface of an NPB and arrowhead indicates nascent rRNA outside NPB. <u>Lower panel:</u> 3D representation of a whole embryo at each stage using Imaris 9.6 for channel segmentation. The hours indicated are “post-injection of hCG”. DNA is labeled in blue and rDNA transcripts in green. *Scale bar = 5µm*. **(B)** Schematic representation of the rDNA sequence and position of the rRNA FISH probes used in this study. Each repeat of the mouse ribosomal sequence (X 82564.1) is transcribed in a long mRNA (47S) encoding for 18S, 5.8S, and 28S rRNAs, separated by two external transcribed spacers (5’ETS and 3’ETS) and two internal transcribed spacer (ITS1 and ITS2). **(C)** Quantification of the rRNA transcription at 2-cell (early (E) and late (L)), 4-cell (e and L), 8-cell (E and L), and 16 cell stages by Real-Time qPCR (RT-qPCR) using primers (described in Table S2) specific of 47S long pre-rRNA, of 5’ETS, ITS1 and ITS2 regions. Four to six biological replicates were done by stage, the expression of these pre-RNA was normalized to a fixed amount of luciferase mRNA added before the reverse transcription. Statistical significance was evaluated using a non-parametric multiple comparison test (nparcomp, R package). ***, p-value < 0.001. **(D)** Localization of H3K9ac (green) and nascent rRNA (magenta) by immuno-RNA-FISH from 2-cell to 8-cell stage. Green arrows indicate H3K9ac bright and big foci close or inside the 5’ETS RNA-FISH signal. Pixels (point by white arrows) that correspond to co-localization between H3K9ac and 5’ETS fluorescent channels were identified using Imaris 9.6 software.

No RNA-FISH signal were detected in the nucleus of 1-cell (data not shown) and early 2-cell stages (36hphCG, Fig 3A, and Fig S2A and B) irrespective of the probe used. At the late 2-cell stage, the 5’ETS signal (corresponding to the nascent transcripts) is located in DAPI-free regions inside the nuclei (arrowhead in Fig 3A, 2-cell embryo at 45hphCG) and at the periphery of bigger NPBs (arrow in Fig 3A). At the 4-cell and 8-cell stages, its localization shifts from the periphery to the inner part of the NPBs. Finally, at the 16-cell stage (75hphCG, Fig 3A), the 5’ETS signal fills the NPBs (Fig 3A - lower panel). As for the 5’ETS probe, the RNA-FISH signal of ITS1 and ITS2 tend to gain space inside the NPBs during progression through early embryonic development (Fig S2 A and B). At the 16-cell stage, ITS2 and 28S as well as ITS1 and 18S are completely intermingled (Fig S2A and B).

To complete RNA-FISH data with quantitative data, real-time quantitative PCR (RT-qPCR) using specific pre-rRNA primers (Table S2) were performed. The level of nascent rRNA transcripts (assessed with 47S and 5’ETS specific primers), gradually increased from late 2-cell to early 8 cell stage and then the curve tends to a plateau (Fig 3C, upper panel). ITS1 and ITS2 specific primers allow us to assess the amount of ‘in-process’ pre-rRNAs: a significant increase was observed from the late 4-cell to the 8-cell stage (Fig 3C, lower panel). Finally, to gain insight into a putative correlation between epigenetic marks and transcription of rDNA, we have determined using an immuno-RNA-FISH approach, the distribution of H3K9ac and H3K4me3 together with their co-localization with nascent transcripts (5’ETS probes). These two histone post-translational modifications (PTMs) are canonically associated with actively transcribed genes (Allis and Jenuwein, 2016). From the 2-cell to the 8-cell stages, we observed that some H3K9ac bright foci (green arrows in Fig 3D) are localized close to 5’ETS RNA-FISH signals (white arrows in Fig 3D). Notably, H3K9ac foci are found at the periphery of 5’ETS signal at 2-cell and 4-cell stages and then dispersed in the 5’ETS cloud inside the NPBs at the 8-cell stage but do not co-localized with the totality of the nascent transcripts (Fig 3D). The same results are found for H3K4me3 (green arrows in Fig S2C).

In conclusion, in contrast to the nucleolus in the differentiated cells, rDNA transcription and rRNA maturation processes are separated in time and space in the immature nucleolus of the embryos at early stages. The first part of the early development (from the late-2-cell to the 4-cell stage) is mainly dedicated to the initiation of rDNA transcription as revealed by a large number of nascent transcripts and the presence of H3K9ac and H3K4me3 localized close to nascent rRNA. The second part of the early development (8-cell and 16-cell stages) is characterized by the maintenance of the initiation of transcription but an increase in the processing of immature pre-rRNAs.

### Inhibition of rRNA transcription and its consequences: elongation is impaired not initiation

To further investigate this temporal separation of initiation of the transcription and processing of rRNA, we used two inhibitors of the RNA Polymerase I (RNA Pol I). Embryos at the 1-cell stage were cultured with either CX-5461 (1µM) or Actinomycin D (ActD, 7.5ng/µL) and fixed 24h (2-cell stage) or 48h later (4-cell stage). We confirmed that both CX-5461-treated and ActD-treated embryos arrested their development at the 4-cell stage (Koné et al., 2016). We then localized the nascent rRNA transcripts by immunoRNA-FISH with 5’ETS probe as well as two key nucleolar components, UBF and Nopp140 involved in the initiation of rRNA transcription and processing of the pre-RNA, respectively (Koné et al., 2016). In both the treated groups, 5’ETS signal (green arrows in Fig 4A), Nopp140 (red arrows in Fig 4A), and UBF (white arrows in Fig 4A) immunoRNA-FISH patterns are altered compared to the control-untreated group (Fig 4A). As described in Koné et al (2016), in the control group of UBF and 5’ETS, signals are superposed and surrounded by Nopp140 signal and in CX-5461-treated embryos, Nopp140 and UBF form nucleolar caps, reviewed in (van Sluis and McStay, 2017) with 5’ETS signals still localized around NPBs. On the contrary, after 48h of treatment, ActD-treated embryos show a higher disruption than CX-5461-treated embryos with complete segregation between the UBF-5’ETS and Nopp140 signals, suggesting that nascent transcripts are not processed anymore.

**Figure 4:**
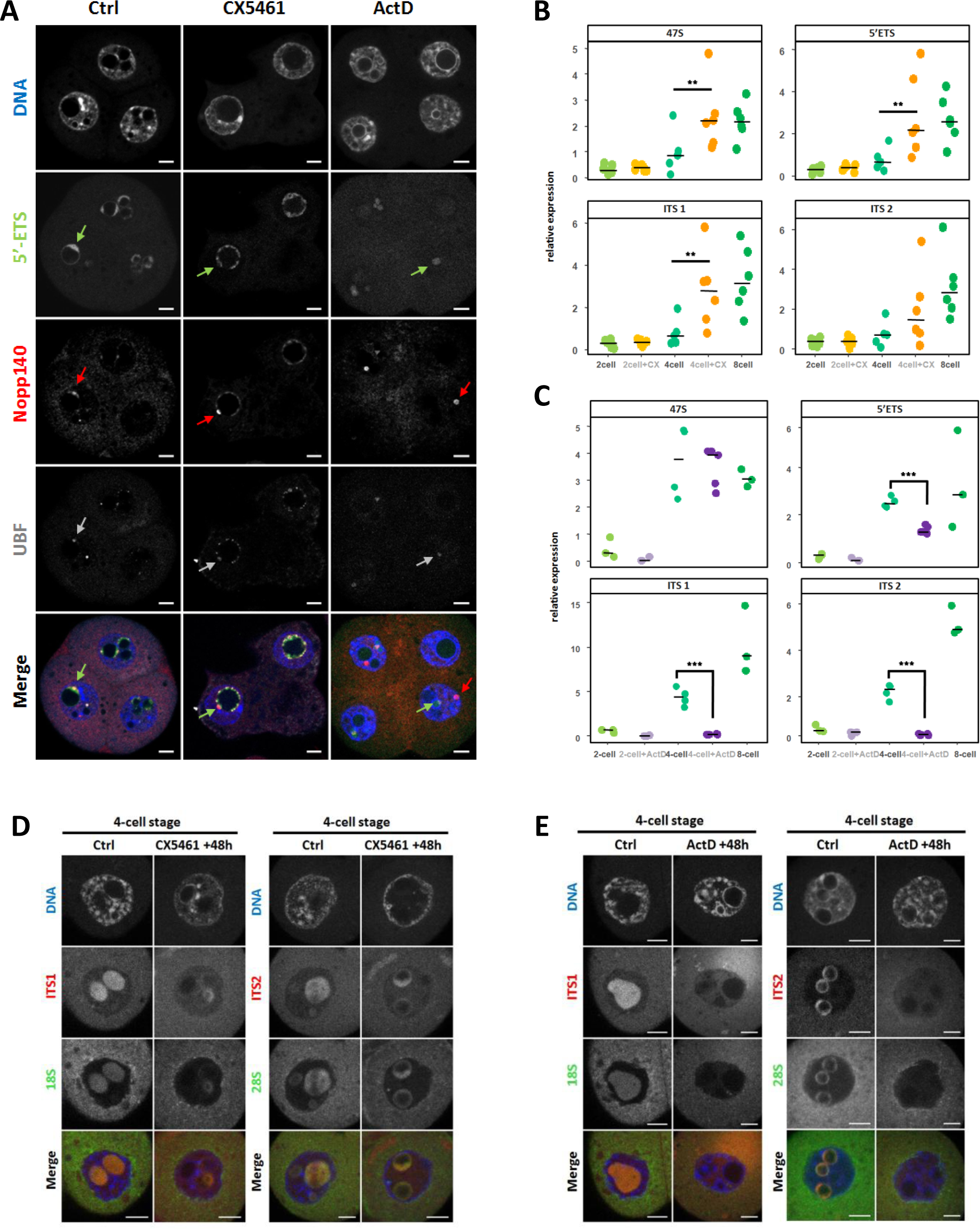
Inhibition of rRNA transcription by CX-5461 and ActD. **(A)** Localization using immuno RNA-FISH of nascent rRNA (5’ETS probe in green), Nopp140 (red) and UBF (grey) at 4-cell stage in non treated (Ctrl) and treated embryos (CX-5461 and ActD), nuclei were counterstained with DAPI (blue). Every single z-section of a confocal stack image show representative nuclei. Green arrows point 5’ETS FISH signals, red arrows Nopp140 immunostaining, and grey arrows UBF immunostaining. *Scale bar = 5µm*. **(B and C)** Quantification by RT-qPCR of the relative expression of rRNAs amplified with specific primers for 47S, 5’ETS, ITS1, and ITS2 in control embryos (2-cell, 4-cell and 8 cell stages, green spots) and treated embryos (2-cell and 4-cell stages): with CX-5461 (B, orange spots) and actinomycin D (C, purple spots). Statistical significance between control and treated embryos was evaluated using the Mann-Whitney test (wilcox, R-package). *, p-value < 0.05, **, p-value < 0.01. **(D and E)** Representative confocal images of 4-cell stages nucleus in non-treated (Ctrl) and treated embryos with CX-5461 (D) or Actinomycin D (E). In process pre-rRNAs species were localized by RNA-FISH with specific fluorescent oligoprobes for ITS1 (red) - 18S (green), left panel or ITS2 (red) - 28S (green), right panel. Nuclei were counterstained with DAPI (blue). *Scale bar = 5µm*.

To test this hypothesis, the number of nascent pre-rRNAs (using primers for 47S-pre rRNA and the 5’ETS regions) and the number of pre-rRNAs in the process (using primers specific to the ITS1 and the ITS2 regions) were assessed by RT-qPCR (Fig 4B and Fig 4C). There was no significant difference between control and treated-embryos at the 2-cell stage. In the CX-5461 condition, the quantity of 47S, 5’ETS, and ITS1 pre-rRNAs were significantly increased (1.8 fold change) between control and the treated 4-cell embryos (Fig 4B, p value< 0.01) although we observed a slight decrease in the RNA-FISH signal compared to control (5’ETS FISH signals in Fig 4A, ITS1 and ITS2 FISH signals in Fig 4D). A putative explanation is that CX-5461-treated embryos compensate a lower number of rDNA loci engaged in transcription by a higher rate of transcription on rDNA that was able to initiate transcription by recruitment of SL1 and UBF. On the contrary, we observed a significant (p-value <0.005) decrease of 5’ETS, ITS1, and ITS2 pre-rRNAs in ActD-treated embryos both by RT-qPCR quantification (Fig 4C) and by RNA-FISH (Fig 4A and 4E). In agreement with the hypothesis that Actinomycin D inhibits RNA Pol I elongation, ITS1 and ITS2 display a stronger reduction (Fig4 E) than 5’ETS (Fig 4A, left panel). CX-5461 has no specific effect on rRNA processing compared to Actinomycin D (in congruance with Mars et al 2020).

Altogether, these results demonstrate that RNA Pol I inhibition with Actinomycin D treatment during early embryonic development induces transcriptional arrest at the 4-cell stage and a massive reorganization of nucleolar proteins due to an incomplete transcription and processing of the long 47S pre-rRNA.

### RNA Pol I inhibition induces changes on the 3D organization of rDNA

To assess the consequences of RNA Pol I inhibition on rDNA organization and the nearby major satellite sequences, DNA-FISH approach was used (Fig. 1). To compare the organization of rDNA and major satellite sequences between the control and treated embryos, the shape of both DNA signals was visualized and analyzed using 3D reconstruction at the 2-cell and 4-cell stages (Fig.5A and 5B) respectively. At the 2-cell stage, the rDNA signals are compacted and juxtaposed to major satellite signals both in the control and treated embryos (green and black arrows in Fig 5A) but their position in regards to NPBs is different between the control (inside NPBs, Ctrl in Fig 5C) and the treated (outside NPBs, CX-5461 and ActD in Fig 5C) embryos, as shown in plot profiles. At the 4-cell stage, the rDNA signal in the treated embryos is less intense than in the control (green arrows in the upper panel, Fig 5B), its spatial distribution is more compact (clusters seem bigger and in lower number) and in ActD-treated embryos, it forms clusters as in 2-cell stage embryos. Plot profiles drawn across the nuclei show that in the CX-5461, rDNA are still inside the NPBs as in the control condition, but not in the ActD condition where rDNA are outside the NPBs, juxtaposed to major satellite sequences (Fig 5D).

**Figure 5:**
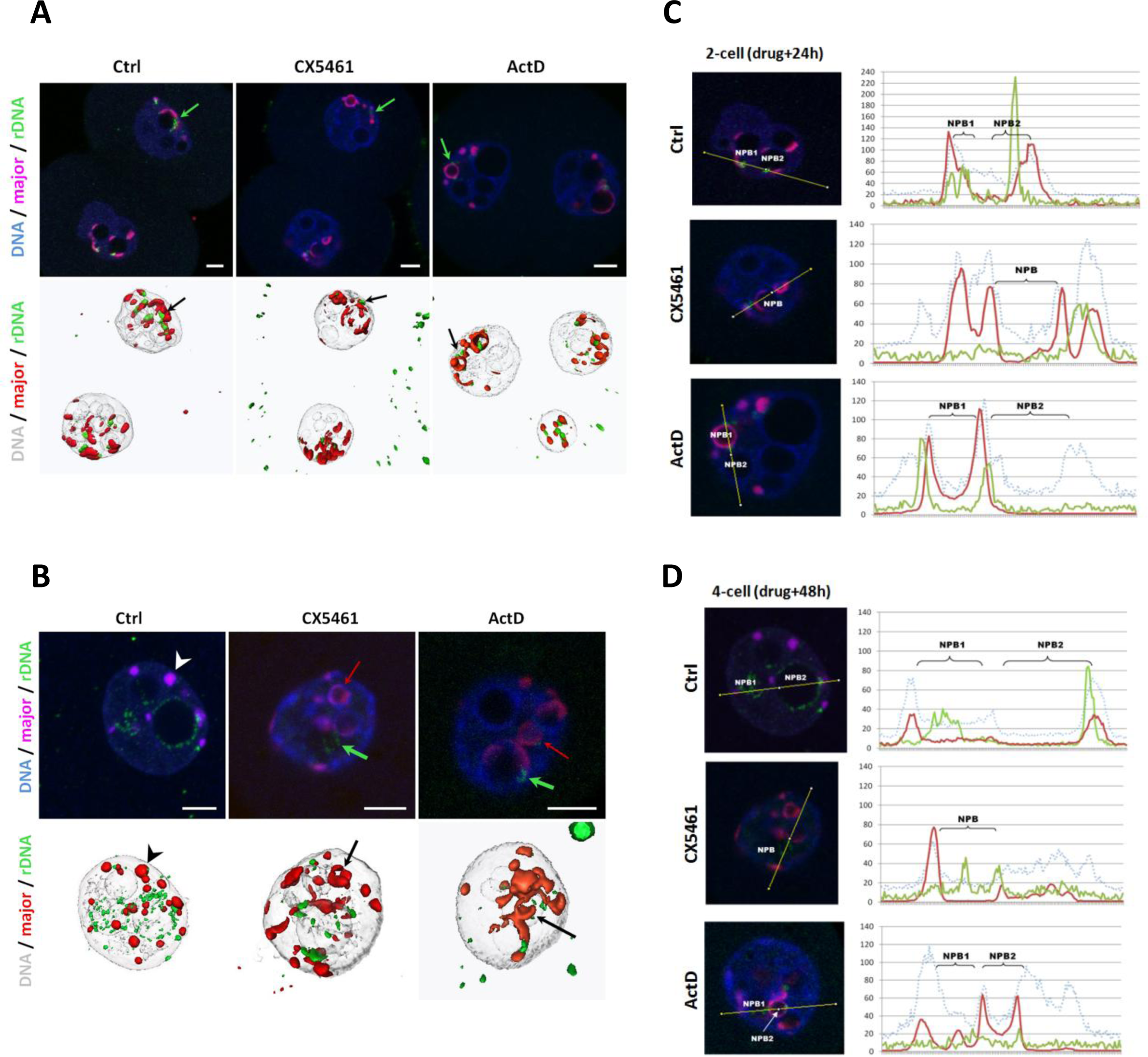
Consequences of RNA Pol I Inhibition on the spatial organization of ribosomal sequences. **(A)** 3D organization of rDNA and major satellite sequences at 2-cell stages detected by 3D DNA-FISH using specific probes. <u>Upper panel:</u> single z-section of representative nuclei of non-treated (Ctrl) and treated embryos (CX-5461 and ActD). <u>Lower panel:</u> Amira 3D representation of 2-cell embryos. Green arrows indicate clustered rDNA FISH signal juxtaposed to major satellite signal at NPBs surface **(B)** 3D organization of rDNA and major satellite sequences at 4-cell stages detected by 3D DNA-FISH. <u>Upper panel:</u> single z-section of representative nuclei of non-treated (Ctrl) and treated embryos (CX-5461 and ActD). <u>Lower panel:</u> Amira 3D representation of each embryo. Green arrows indicate clustered rDNA FISH signal and red arrows major satellite sequences FISH signal with a ring/half-ring shape. White arrowhead point major satellite sequences FISH signal with a round shape called chromocenter. **(C and D)** Localization of rDNA and major satellite signals in regards to NPBs boundaries at 2-cell stages (C) and 4-cell stages (D) in non-treated (Ctlr) and treated (CX-5461 or ActD) embryos. <u>Left panel:</u> single z-section of a representative nucleus with the line used to draw with Fiji the Intensity plot profile. <u>Right panel:</u> Fluorescence intensity plot profile along the line across multiple channels (blue dot line for DAPI, the green line for rDNA signal, and red line for major satellite signal). DNA is in blue or grey, rDNA in green, and major satellite sequences in magenta. *Scale bar = 5µm*. mPN: major : mouse major satellite sequences, rDNA: ribosomal DNA.

To assess and quantify the differences in terms of compaction and shape between the control and the treated embryos, rDNA FISH signals were segmented using Imaris software (as previously in Fig 2B and 2C) to measure the volume and the sphericity of the rDNA signal at 2-cell and 4-cell stages (Fig 6A and 6B). In the controlled condition, the mean volume of the rDNA signal increases from the 2-cell to 4-cell stage, and its shape (measured by the sphericity) becomes less round meaning that rDNA sequences decondensed concomitantly with being actively transcribed. At the 2-cell stage, the shape of the rDNA signal is significantly different between the control and the treated embryos (left panel in Fig 6B, p-value <0.005) suggesting that ribosomal sequences are less round in the treated embryos and putatively less compact. At the 4-cell stage, the mean volume of rDNA signal is significantly (p-value <0.005) different between the control and the treated embryos (Fig 6B, 5.31 +/-1.63 in Ctrl vs 0.85+/-0.48 and 0.93+/- 0.27 in CX-5461 and ActD treated embryos, respectively) meaning that rDNA sequences stay tightly compact and clustered upon RNA Pol I inhibition.

**Figure 6:**
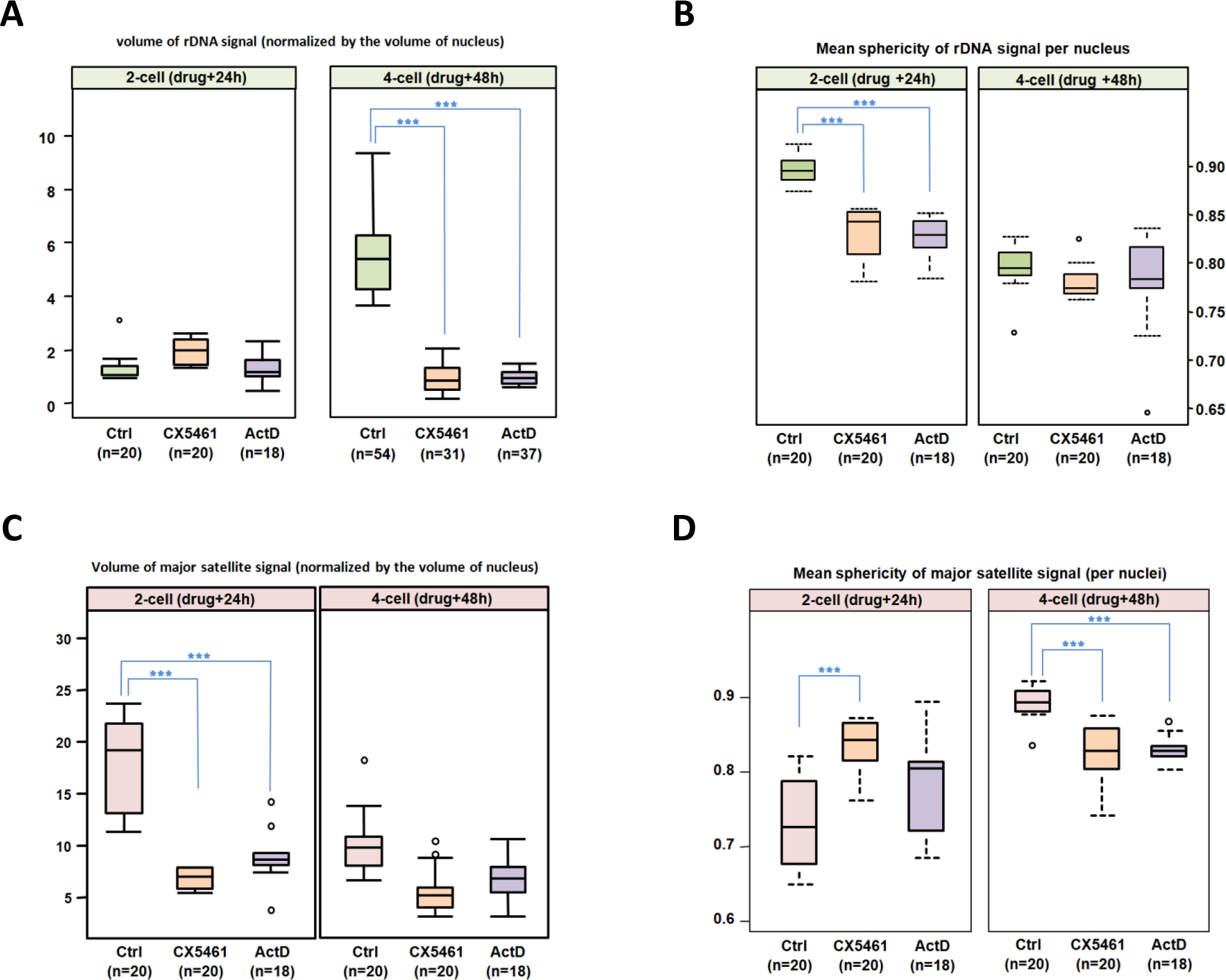
Consequences of RNA Pol I Inhibition on rDNA and major satellite sequences 3D conformation (volume and shape). **(A and B)** Quantification of rDNA total volume (A) and average sphericity (B) per nucleus and stage based on the DNA-FISH signal of the ribosomal sequences at 2-cell and 4-cell stages in non-treated (Ctrl) and treated (CX-5461 or ActD) embryos. The number of nuclei examined per condition is indicated below the name of the condition (Ctrl in green, CX-5461 in orange, and ActD in purple). **(C and D)** Quantification of major satellite sequence total volume (C) and average sphericity (D) per nucleus and per stage based on the DNA-FISH signal, at 2-cell and 4-cell stages in non-treated (Ctrl) and treated (CX-5461 or ActD) embryos. The number of nuclei examined per condition is indicated below the name of the condition (Ctrl in pink, CX-5461 in orange, and ActD in purple). All measurements were done using Imaris 9.6 (Oxford Instruments). Statistical significance was evaluated using a non-parametric multiple comparison test (nparcomp, R package). ***, p-value < 0.001.

### Disruption of 3D organization of rDNA with RNA Pol I inhibition has consequences on the transcription of pericentromeric sequences

By paying attention to the major satellite signals (shape and volume) assisted us in pinpointing that their distribution is modified by the drug treatment. Indeed, after 48h of RNA Pol I inhibition, major satellite DNA-FISH signals show a croissant shape- as in 2-cell embryos (red and black arrows in Fig 5B)- whereas they are normally clustered into chromocenters at the 4-cell stage in controls (black arrowhead in Fig 5B). To confirm this further, the mean volume and the sphericity of the DNA-FISH signals of major satellite sequences were measured using Imaris in the control and the treated embryos (Fig S3A and S3B and Fig 6C and 6D). The mean volume of major satellite sequences FISH signals is significantly lower (p-value <0.005) at the 2-cell stage in the treated embryos compared to the control (right panel, Fig 6C). At the 4cell stage, the volume in the control decreases, while there is no change in the treated embryos (left panel, Fig 6C). Furthermore, the decreased sphericity in the treated embryos compared to the control at the 4-cell stage (right panel, Fig 6D) means that the major satellite sequences are less round and compact in the treated embryos. Similarly, the mean sphericity of the 4-cell stage in the treated embryos is not significantly different from that of the 2-cell control embryos.

Previous studies highlighted that major satellite sequences are specifically transcribed at the 2-cell stage when their shape is less round (Casanova et al., 2013; Probst et al., 2010), therefore we decided to assess major satellite transcription in the treated vs. control embryos by RNA-FISH and RT-qPCR. No difference in the major satellite RNAs was detected between the control and treated embryos at the 2-cell stage, neither by RNA-FISH nor by RT-qPCR (Fig 7A, B and C) however we detected a significant decrease of transcription in the 4-cell stage ActD-treated embryos (Fig 7B, p< 0.005). On the other hand, a housekeeping gene such as *Hprt* (Fig S3C) does not vary in ActD-treated embryos suggesting a reciprocal influence between rDNA and major satellite transcription. As a matter of fact, after the 2-cell stage burst of transcription, the remaining major satellite RNA-FISH spots were always visualized inside the cloud of 5’ETS FISH signal (green arrows in Fig 7D), at the 4-cell, 8-cell, and 16-cell stages (red arrow in Fig 7D) suggesting that rDNA transcription might induce a chromatin environment that favors transcription of major satellite sequences.

**Figure 7:**
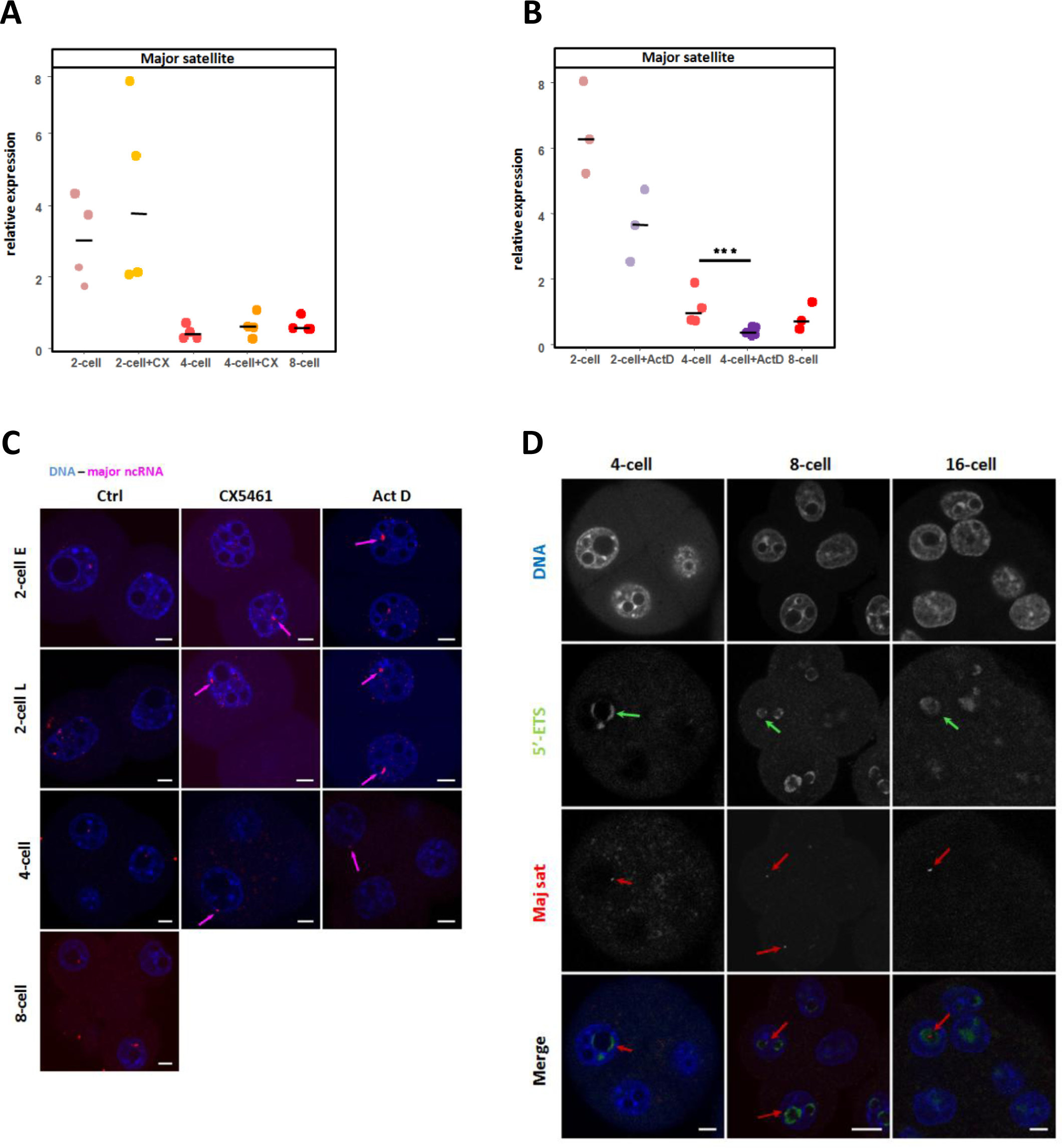
Consequences of RNA Pol I Inhibition on the expression of major satellite sequences. **(A and B)** Quantification by RT-qPCR of the relative expression of ncRNA amplified with specific primers for major satellite sequence in control embryos (2-cell, 4-cell and 8 cell stages, pink/red spots) and treated embryos (2-cell and 4-cell stages): with CX-5461 (A, orange spots) and actinomycin D (B, purple spots). Three to four biological replicates were done by condition, the expression of these ncRNA was normalized to a fixed amount of luciferase mRNA added before the reverse transcription. Statistical significance between control and treated embryos was evaluated using the Mann-Whitney test (Wilcox, R-package). **, p-value < 0.01. **(C)** Major satellite transcripts (magenta) visualized using RNA-FISH in non treated (Ctlr) in non-treated (Ctrl) and treated (CX-5461 or ActD) embryos from 2-cell (early (E) and late (L)) to 4-cell and 8-cell stages. ncRNAs spots are indicated by red arrows in treated embryos. **(D)** Localization of nascent rRNAs (green) and major satellite ncRNAs (red) in mouse embryos from the 4-cell to 16-cell stages by RNA-FISH using specific probes for 5’ETS region and major satellite sequences. Green arrows indicate a 5’ETS RNA-FISH signal closed to a major satellite signal (red arrows). Nuclei were counterstained with DAPI (blue). *Scale bar = 5µm*.

## Discussion

### Spatial organization of rDNA during early development

In this study, we demonstrated that the spatial positioning of ribosomal genes (rDNA) correlates with their transcription activity. In the context of early embryonic development, ribosomal repeat sequences are first clustered at the surface of NPBs, flanked by ring-shaped major satellite sequences (1-cell and early 2-cell). From the late 2-cell to the 16-cell/morula stage, the sequences then expanded in the inner space of the NPBs until complete filling. During this period, rDNA gradually separated spatially from major satellite sequences although they are physically attached on the same chromosome. The cloudy organization of the active ribosomal sequences we observed fits with the localization of UBF as previously described (Fulka and Langerova, 2014; Hamdane et al., 2016; Koné et al., 2016) and is specific to the early embryonic period. This unique nucleolus organization is very different from the somatic ones (Maiser et al., 2020) that can only be found at the morula and blastocyst stages (Lavrentyeva et al., 2015). Interestingly, this second switch in the nucleolus conformation is associated with a second burst of embryonic genes transcription at the morula-to-blastocyst transition described in transcriptomic data (Hamatani et al., 2004; Hamatani et al., 2006; Wang et al., 2004).

In parallel to these drastic changes in the spatial organization of the sequences, we show that rRNA synthesis begins at the mid-2-cell stage, first in the small spherical structures outside the NPBs. Then, nascent rRNAs later appear at the surface of some NPBs as previously described (Koné et al., 2016; Lavrentyeva et al., 2015). Therefore, at very early stages (1-cell and 2-cell stages), NPBs can not be considered as a reservoir for pre-rRNAs waiting to be cleaved confirming previous studies (Lavrentyeva et al., 2015; Shishova et al., 2015a; Shishova et al., 2015b). Interestingly, immuno-RNA-FISH revealed that bright spots of H3K9ac and H3K4me3 are located at the vicinity of 5’ETS signal at the 2-cell stage and that these spots are found inside the 5’ETS cloud at later stages. These results are in agreement with ChIP data obtained on mouse embryonic fibroblast or 3T3 cells (Herdman et al., 2017; Zentner et al., 2014) describing an enrichment of these two epigenetic marks on the spacer promoter (UCE core) at the actively transcribed rDNA repeat (Grummt and Längst, 2013; Moss et al., 2019).

### Inhibition of RNA Pol I and its impact on rRNA processing

To decipher the link between the spatial organization of rDNA and their transcriptional activity, we used two drugs that inhibit RNA Pol I activity in two different mechanisms. CX-5461 is a molecule that inhibits the initiation of transcription by impairing the formation of the pre-initiation complex (PIC) (Drygin et al., 2011) whereas Actinomycin D is a DNA intercalant that inhibits elongation by the RNA polymerases (all of them) in a dose-dependent manner (Perry and Kelley, 1970). Several recent studies (Bruno et al., 2020; Mars et al., 2020; Sanij et al., 2020) discussed the specific targets of CX-5461 (SL1 or Topoisomerase II or RRN3) and its mode of action (G-quadruplex stabilization, DNA damage, impairing of DNA replication fork). Notably, Mars et al., (2020) emphasis that RNA Pol I inhibition by CX-5461 is not reversible and can lead to cell death because of DNA damage. To get read of these putative side effects of CX-5461, ActD was also used in this study.

In the early embryonic context, we observed that CX-5461 disrupts the classical tripartite (i.e. FC, DFC and Granular component (GC) (Boisvert et al., 2007; Pederson, 2011)) nucleolus organization (this study & Koné et al 2016). In agreement with (Mars et al., 2020), our study demonstrates that CX-5461 has less impact on the elongation process than Actinomycin D. In the treated embryos, Actinomycin D treatment also inhibits nascent rRNA synthesis but induces a complete stop of the long pre-rRNA production (labeled with ITS1 and/or ITS2) which in turn induces both dissociation of transcription initiation (labeled by 5’ETS and UBF) and processing of rRNA (labeled with Nopp140). It is worth-mentioning that the treated embryos arrested their development at the 4-cell stage most likely because maternally inherited ribosomes are not renewed as demonstrated in the embryos invalidate for UBF (Hamdane et al., 2014; Hamdane et al., 2016).

### The link between major satellite and rDNA sequences transcription

Lastly, we compared rDNA shape, distribution, and transcriptional activity in time and space with regards to major satellite sequences during early development. Indeed, major satellite sequences are drastically reorganized at the time of EGA (Aguirre-Lavin et al., 2012; Probst et al., 2007) and this reorganization is dependent on major satellite transcripts (Casanova et al., 2013; Probst et al., 2010). On the other hand, localization of major satellite sequences around the NPBs is mandatory at the 1-cell stage as demonstrated by enucleolation experiments (Fulka and Langerova, 2014; Kyogoku et al., 2014; Ogushi and Saitou, 2010; Ogushi et al., 2008) and at the early 2-cell stages (Jachowicz et al., 2013) even if in-depth knowledge of the mechanisms involved are still unknown. Growing pieces of evidence propose that NPBs, rather than acting as a precursor of nucleoli, act as a platform or a Velcro for major satellite sequences (Fulka and Aoki, 2016; Padeken and Heun, 2014) thanks to nucleoplasmin 2 (NPM2, (Burns et al., 2003; Inoue and Aoki, 2010; Kyogoku et al., 2014)). In fact injection of sole NPM2 mRNA rescue developmental failure of enucleolated oocytes (Ogushi et al., 2017). Surprisingly, the absence of NPBs induces the earlier formation of chromocenter-like structures at the zygotic stage (Ogushi et al., 2017) although transcription of major satellite sequences is repressed after the end of the 2-cell stage (Fulka and Langerova, 2014). In this study, we show that inhibition of rDNA transcription induces major satellite sequences to change from a chromocenter-like shape to a ring-like shape without increasing their transcription activity. Surprisingly, Actinomycin D treatment induces a decrease of major satellite transcription while other RNA polymarase II-dependent genes are not affected.

rDNA spatial organization is disrupted on the inhibition of RNA Pol I and rRNA sequences aggregate, exhibiting a round shape comparable to earlier inactive stages. Similarly, major satellite sequences exhibit a spatial organization that looks like an early 2-cell stage. This loss of chromatin organization in RNA Pol I inhibited embryos can partially explain the decrease of major satellite sequences transcription. After the 2-cell stage, major satellite transcription is severely repressed and only one or two spots of major satellite RNA can be detected per nucleus by RNA-FISH (Probst et al., 2010). Remarkably, these spots were often found inside the 5’ETS cloud (this study), suggesting transcription of major satellite sequences after the 2-cell stage may require a suitable environment that favor transcription. A putative explanation -among others-is that transcription of major satellite sequences, in 4-cell and 8-cell embryos, need the rDNA peculiar chromatin state (Moss et al., 2019; Potapova and Gerton, 2019): nucleosome-free chromatin with UBF that promotes loops formation (Stefanovsky et al., 2001).

## Material and Methods

### Ethics

Animal care and handling were carried out according to the national rules on ethics and animal welfare in the Animal facility (IERP, INRAE, Infectiology of fishes and rodent facility, doi: 10.15454/1.5572427140471238E12, Jouy-en-Josas, C78-720). This work was approved by the French Ministry of Higher Education, Research and Innovation (n°15-55) and the local ethical committee (INRAE Jouy-en-Josas Center). Departmental veterinary regulatory services have delivered habilitation to work with laboratory animals to ABG (n°78–184) who supervised the work.

### Collection of mouse embryos and culture

Embryos were collected upon superovulation and mating of C57/CBAF1 mice as previously described (Koné et al., 2016). Zygotes were obtained by dissecting the ampulla of the oviduct 24 hours post-injection of human chorionic gonadotropin (hCG) and treated briefly with 1mg/ml of hyaluronidase (Sigma) in M2 medium (Sigma) to remove follicular cells. Then, 1-cell (24h phCG) embryos were cultured *in vitro* in M16 medium (Sigma) at 37°C in a humidified atmosphere of 5% CO_2_ until they reached the adequate stages to be processed: early 2-cell (36h phCG - 2CE), late 2-cell (48h phCG - 2CL), early 4-cell (54h phCG - 4CE), late 4-cell (58h phCG - 4CL), early 8-cell (60h phCG - 8CE), late 8-cell (72h phCG - 8CL), 16-cell (75h phCG), Morula (96h phCG) and Blastocyst (110h phCG).

### CX-5461 and Actinomycin D treatment

Two different RNA Polymerase I (RNA Pol I) inhibitors were used in this study: CX-5461 that impairs the formation of the RNA Polymerase I pre-initiation complex (PIC) by disrupting the recruitment of SL1 (Selective factor 1) with UBF (Upstream biding factor) to the rDNA promoter (Drygin et al., 2011) and Actinomycin D that intercalates into the DNA and thereby inhibits elongation of all the RNA polymerases activity, although with a higher affinity for RNA Pol I (Bensaude, 2011; Perry and Kelley, 1970). These two drugs induce segregation of the nucleolus components dedicated either to transcription (fibrillar and dense fibrillar components -FC and DFC) or to rRNA processing (granular component, GC). These components can be visualized by fluorescent immuno-detection of the proteins involved in these processes (UBF and Nopp140 respectively, as described in (Koné et al., 2016)). Embryos collected at the 1-cell stage (24hphCG) were cultured in M16 medium containing either 1µM of CX-5461 (Adooq) or 7.5ng/µL of Actinomycin D (Sigma) to inhibit RNA Pol I as described respectively in (Bellier, 1997; Koné et al., 2016). Every 24h, the embryos were transferred into new droplets of medium containing the RNA Pol I inhibitors. The treated embryos were fixed at two-time points: 24h and 48h after exposure to the drug and then processed for immunoRNA-FISH / DNA-FISH or frozen dry in low RNA binding tubes and kept at -80°C for RT-qPCR assays.

### Probes for DNA and RNA-FISH

rDNA and major satellite FISH probes were prepared as described in (Aguirre-Lavin et al., 2012) and (Bonnet-Garnier et al., 2013). Briefly, cosmid containing the entire transcribed sequences (n°13, detailed in (Akhmanova et al., 2000; van de Nobelen et al., 2010) was used for the ribosomal sequences (kind gift from N. Galjart). The amplified rDNA gene fragments were purified using nucleobond AX 100 columns for Miniprep System (Macherey-Nagel, ref 740521.100). The rDNA sequences were then labeled with Digoxigenin-11-dUTP by nick translation according to the manufacturer’s protocol (Roche, Ref 11277065910). For the detection of major satellites, a probe prepared by PCR on genomic mouse DNA with the primers 5’-CATATTCCAGGTCCTTCAGTGTGC-3’ and 5’-CACTTTAGGACGTGAAAT ATGGCG-3’ was labeled with Cy3 by random priming according to the kit instruction (Invitrogen Kit, Ref 18095–011).

Specific oligo-probes (designed from the complete 45S mouse ribosomal sequences - X82564.1 or adapted from (Le Bouteiller et al., 2013; Shishova and others, 2011; Shishova et al., 2015b) directly labeled with a fluorophore in 3’ end (sequences are detailed in Table S1) were purchased from Eurogentec. These probes located (Fig. 2B) on different regions of the 47S pre-RNA (Kent et al., 2008) are named: 5’ETS for the probe specific to the external transcribed spacer (rapidly removed, allowing the detection of early transcription sites, i.e. nascent rRNA transcripts), ITS1: specific for the internal transcribed spacer 1 located between the 18S and the 5.8S regions and ITS2: specific for the internal transcribed spacer 2 located between the 5.8S and the 28S region. ITS1 and ITS2 enable the mapping of the region where rRNA processing occurs. Probes specific for 18S and 28S regions were also used to assess if mature or pre-rRNAs were already accumulated in NPBs before the onset of rDNA transcription. Of note, the 18S and 28S probes signal can be detected in the cytoplasm because they are components of ribosomes.

### DNA-FISH

For DNA-FISH, DNA probes described above specific to either ribosomal or major satellite sequences were used together on the whole embryos. During all the manipulations, drying was avoided to preserve the 3D of nuclei. This protocol is based on (Miyanari and Torres-Padilla, 2012) with slight modifications. The embryos collected at a specific stage were transferred in a 10-well glass dish warm at 37°C. The embryos were rinsed in M2 medium, submitted briefly (less than 15s) to Tyrode’s acid solution (Sigma-Aldrich), washed quickly in an M2 with 10mM phenylmethanesulfonyl fluoride (PMSF, Fluka) solution then in a 0.5% polyvinyl-pyrrolidone (PVP) in PBS with 10mM PMSF solution and finally incubated in a fixation/permeabilization solution (4% paraformaldehyde (PFA), 0.5% Triton X-100, 10mM PMSF, 0.5% PVP in PBS) for 15min at 37°C. The embryos were then washed twice in a PBS/PVP 0.5% solution at RT and let at +4°C (at least overnight) in a humid chamber until fixation of other stages. To remove the zona pellucida (ZP), all the embryos were incubated under a stereomicroscope to monitor the ZP vanishing in a 0.1N HCl (Prolabo) solution for 20 to 30s (depending on the stage) at RT and transferred to PBS/PVP 0.5%. The ZP removal was achieved by successive passages through a very fine glass pipette when needed. The embryos were permeabilized in a 0.5% Triton X-100 / 200µg/ml RNase solution for 30 min at 37°C and washed twice in 2X saline-sodium citrate (2XSSC) pH 6.3 for 5 min at 37°C. The embryos were then transferred for 3h at 37°C in a humidified chamber in a 20µl drop of hybridization buffer (50% formamide, SCC 2X, Denhardt 1X, 40 mM NaH2PO4, 10% dextran sulfate) containing the DNA-FISH probes mix (14µl of rDNA probe solution at 50ng/µl and 1µl of major satellite probe solution at 100 ng/µl completed to 20µl with hybridization buffer). The embryos were denatured at 85°C for 10 min on a heating plate and then let three days for hybridization at 37°C in a humid chamber. After hybridization, the embryos were rinsed twice with 2X SSC / 0.1% Triton X-100 / 0.5% PVP for 10 min at 42°C. Embryos were then incubated in a blocking solution (4X SSC / 2% BSA) for 1h at RT and incubated with a primary antibody diluted in blocking solution (sheep anti-digoxigenin, 1/200, Roche) overnight at +4°C in a dark humidified chamber. The embryos were washed twice in 4X SSC / 2% BSA / 0.05% Tween20 before and after incubation with the secondary antibody (mouse anti-sheep Ig coupled to FITC, 1/200, 713-095-147, Jackson ImmunoResearch Laboratories Inc., USA) for 1h at RT. The embryos were washed in 2X SSC / 0.5% PVP and gently mounted to preserve the 3D structure of the nuclei between slides (Menzel Superfrost Plus, Thermo Scientific) and cover-slip with a large amount of Vectashield antifading agent (Abcys) containing 10µg/mL DAPI (Invitrogen).

### RNA-FISH

For RNA-FISH, the embryos were processed almost as for DNA-FISH with two main differences: 1) the ZP was not removed by HCl treatment; 2) an RNAse inhibitor (1µL/mL of RNasin - Promega) was added to all the solutions after the fixation/permeabilization step. Briefly, after fixation/permeabilization, embryos were further permeabilized in 0.5% Triton X100 /0.5% PVP-PBS for 45min at 37°C, washed and transferred for 30min at 50°C in the pre-hybridization mix containing 50% Formamide (Sigma-Aldrich); 0.5µg/µL tRNA (Sigma-Aldrich); 1X hybridization buffer (2X hybridization buffer was prepared beforehand with 20% dextran sulfate, 4X SSC; 1mM EDTA, 40mg/mL BSA, 2mg/mL PVP, 0.1% Triton X100, diluted in pure grade water). In parallel, a solution containing the fluorescent oligonucleotide probes (100ng/µl diluted in 1X hybridization buffer) was denatured 10 min at 85°C and immediately put on ice to avoid renaturation of the DNA. The embryos were then transferred in this hybridization mix and incubated overnight at 42°C. The embryos were washed twice in 2XSSC; 0.5%PVP; 0.1% Triton X100 (diluted with pure grade water) for 10min at 37°C and mounted on slides with coverslips as described in the DNA-FISH section.

### ImmunoRNA-FISH

Immunofluorescence labeling followed by RNA-FISH was performed as described in Kone et al. (2016) with slight modifications. Briefly, the embryos were fixed as described in the RNA-FISH section before immunostaining. The embryos were then permeabilized in 0.5% Triton X100 / 0.5% PVP-PBS for 30min at 37°C and transferred in 2% BSA-PBS 1h at RT. The embryos were incubated with primary antibody overnight at 4°C, washed thrice in 0.5%PVP-PBS, and then incubated with secondary antibody for 1h and post-fixed in 2% PFA/0.5% PVP-PBS for 10 min at RT. The embryos were then further processed for RNA-FISH as described in the RNA-FISH section. All antibodies were diluted in 2% BSA-PBS. Again, RNAse inhibitor (1µL/mL of RNasin - Promega) was added to all the solutions after the fixation/permeabilization step.

### Antibodies

Primary antibodies used: anti-UBF (1/100; mouse polyclonal antibody H00007343-M01; Novus Biologicals), anti-Nopp140 (1/150; rabbit polyclonal antibody, RF12 serum; a gift from U. Thomas Meier, Department of Anatomy and Structural Biology, New York, USA), anti-H3K9ac (1/100, rabbit polyclonal antibody, 39917) and anti-H3K4me3 (1/400, rabbit polyclonal antibody, ab8550, Abcam). Secondary antibodies used: IgG donkey anti-mouse Cy3 (1/200, 715-025-151, Jackson ImmunoResarch) and IgG donkey anti-rabbit Cy5 (1/200, 111-175-152, Jackson ImmunoResarch).

### Confocal microscopy and image analyses

Imaging was performed with a ZEISS LSM 700 confocal laser scanning microscope (MIMA2, INRAE, Microscopy and Imaging Facility for Microbes, Animals and Foods, https://doi.org/10.15454/1.5572348210007727E12) equipped with a 63X (1.4 NA) oil immersion objective. Z-stacks were acquired with a frame size of 512 × 512 or 1024 x 1024, a pixel depth of 8 bits, and a z-distance of 0.37 µm between optical sections. Fluorescence wavelengths of 405, 488, 555, and 639 nm were used to excite DAPI, Alexa-488, or FITC, Cy3, and Cy5, respectively.

Fluorescent profiles measurement were generated in Fiji (Schindelin et al., 2012; Schindelin et al., 2015; Schneider et al., 2012) and 3D reconstructions using AMIRA 3.1 software (Mercury Computer Systems, Berlin, Germany, (Stalling et al.)

Quantitative image analysis was performed manually using Imaris 9.6 software (Oxford Instruments) available at the MIMA2 ISC INRAE. Briefly, in 10 embryos of each stage, nuclei were first segmented using an automatically set threshold (wizard function in Imaris surfaces package). Major satellites and rDNA signals were then segmented in a specified region of interest (ROI defined by the nucleus label) following the same process. For all detected objects, several values (area, volume, sphericity, numbers) were calculated by Imaris and exported for statistical analysis in Microsoft Excel file format. For each analyzed nuclei, the sum of the volume of DNA-FISH signals was calculated to obtain the total volume of rDNA or major satellite DNA-FISH signals in a given nucleus. Then this total volume was normalized by the volume of the given nucleus using this formula: (Vol_tot_FISH_ / Vol_tot_nucleus_)x 100, to circumvent any nucleus size effect when comparing between stages.

### RT-qPCR

The embryos at each stage were pooled by a batch of 10 and snap-frozen. Total RNA was prepared without purification using the SingleShot Cell Lysis kit (Biorad) that includes DNase treatment. cDNA was prepared directly from the lysates with random hexamers (300 ng) (Lifetechnologies, France) following the supplier’s protocol (25°C for 5 min, 50°C for 60 min and 70°C for 15 min) in the presence of 10 fg of Luciferase RNA (Promega) using the SuperSript III reverse transcription kit (Invitrogen). Negative controls without reverse transcriptase were prepared the same way. RT-qPCR was performed on 0.2 or 0.032 embryo equivalent (as described in Salvaing et al 2016), depending on the genes, using KAPA SYBR FAST (Roche) or SYBR Green (Applied, for Major satellites) master mix on a StepOne Plus cycling machine (Applied). The difference in amplification of positive and negative samples was at least 5 Ct (5 to 9). To avoid biais due to a highly variable amount of transcript from endogenous genes (Hprt1) before the EGA, the results were finally normalized to Luciferase (as described in Bui et al). Primers are listed in Table S2. Three to six biological replicates were used.

### Statistical analyses

All the statistical analyses and tests were performed using R packages. Several R packages such as ggplot2 and lattice were used in complement to Rcmdr packages to generate graphs. The normality and homogeneity of variances were tested using the ’shapiro.test’ and ’bartlett.test’ R-packages, respectively. As the sample size was small (<30), non-parametrical tests were used. Two by two comparison were done using ’wilcox’ packge (Mann-Whitney-Wilcoxon) and comparison of the two consecutive stages were done with ’nparcomp’ package (Nonparametric Multiple Comparisons).

### Supplementary material

Figs. S1–S5 show 3D organization of ribosomal sequences (rDNA) at 1-cell stage, 16-cell and morula to blastocyst stages using DNA-FISH, the number of nucleolar precursor bodies (NPBs), Quantification of the volume of nucleus and rDNA FISH signals using Imaris, Localization by RNA-FISH of several ribosomal RNA (rRNA) from 2-cell to 16-cell stages, Localization of H3K4me3 and nacent transcripts of rRNA by immunoRNA-FISH, Quantification of major satellite sequences volume and sphericity from 2-cell up to 16-cell stage and relative expression of an housekeeping gene (*Hprt*) by RT-qPCR in control and treated embryos.

Tables S1–S2 show the list of oligoprobes used for the RNA-FISH and the list of primers used in th RT-qPCR experiments respectively.

## Abbreviations

5’ETS: 5’ externally transcribed spacer
ActD: Actinomycin D
EGA: Embryonic Genome Activation
hphCG: hours post-injection of human Chorionic Gonadotrophin
H3K9ac: Histone 3 acetylated in lysine 9
H3K4me3: Histone 3 tri-methylated in lysine 4
ITS1: internal transcribed spacer 1
ITS2: internal transcribed spacer 2
Major sat: mouse major satellite sequences
NPB: Nucleolar Precursor Body
Nopp140: Nucleolar phosphoprotein of 140 kD
mPN: maternal pronucleus
pPN: paternal pronucleus
PVP: Poly Vinyl Pyrrolidone
RT-qPCR: Real-Time quantitative PCR
RNA polymerase I: RNA pol I
rDNA: ribosomal DNA genes/sequences
rRNA: ribosomal RNA
UBF: Upstream Binding Factor

## Acknowledgements

We would like to thank Prof. Galjart Niels (Nederland), M Cohen-Tanoudji (Pasteur Institute), and Janice Britton Davidian lab (Montpellier) for their kind gifts of rDNA plasmids/cosmids and the RNA-FISH probes. We also thank Claire Boulesteix for her technical assistance and laboratory facilities and Bertrand Bed’hom for his help with Imaris analysis. We thank the ISC MIMA2 (Microscopy and Imaging Facility for Microbes, Animals and Foods, doi: 10.15454/1.5572348210007727E12) and particularly Pierre Adenot for advice in 3D images analyses with ImageJ and Imaris. We acknowledge the staff of the INRAE Infectiology of Fishes and Rodents Facility (IERP-UE907, Jouy-en-Josas Research Center, France) in which animal experiments have been performed. IERP Facility belongs to the National Distributed Research Infrastructure for the Control of Animal and Zoonotic Emerging Infectious Diseases through In Vivo Investigation (EMERG’IN DOI: 10.15454/1.5572352821559333E12).

This project was funded by the REVIVE Labex (Investissement d’Avenir, ANR-10-LABX-73) and supported by the PHASE Department of the French National Research Institute for Agriculture, Food and Environment (INRAE).

The authors declare no competing financial interests.

## Authors contribution

M. Chebrout set up the RNA-FISH and immunoRNA-FISH experiments, performed majority of the experiments, and analyzed the data. M.C. Koné, M. Cournut and H.U Jan performed and analyzed DNA-FISH experiments. R. Fleurot and T. Aguirre-Lavin set up and performed the DNA-FISH experiments. A. Jouneau and N. Peynot performed and analyzed the RT-qPCR experiments. N. Beaujean secured funding, gave guidance during initial data production and analysis, and made early corrections to the manuscript. A. Bonnet-Garnier supervised, directed, and designed the study, analyzed, and interpreted the data, made the figures, and wrote the article. All the authors reviewed and commented on the manuscript.

## Ethical approval

All applicable international, national, and/or institutional guidelines for the care and use of animals were followed. This article does not contain any studies with human participants performed by any of the authors.

## References

Adenot, P. G., Mercier, Y., Renard, J.-P. and Thompson, E. M. (1997). Differential H4 acetylation of paternal and maternal chromatin precedes DNA replication and differential transcriptional activity in pronuclei of 1-cell mouse embryos. Development 124, 4615–4625.

Aguirre-Lavin, T., Adenot, P., Bonnet-Garnier, A., Lehmann, G., Fleurot, R., Boulesteix, C., Debey, P. and Beaujean, N. (2012). 3D-FISH analysis of embryonic nuclei in mouse highlights several abrupt changes of nuclear organization during preimplantation development. BMC Dev. Biol. 12, 30.

Akhmanova, A., Verkerk, T., Langeveld, A., Grosveld, F. and Galjart, N. (2000). Characterisation of transcriptionally active and inactive chromatin domains in neurons. J. Cell Sci. 113, 4463– 4474.

Allis, C. D. and Jenuwein, T. (2016). The molecular hallmarks of epigenetic control. Nat. Rev. Genet.

Baran, V., Veselá, J., Rehák, P., Koppel, J. and Fléchon, J. E. (1995). Localization of fibrillarin and nucleolin in nucleoli of mouse preimplantation embryos. Mol Reprod Dev 40,.

Baran, V., Brochard, V., Renard, J. P. and Flechon, J. E. (2001). Nopp 140 involvement in nucleologenesis of mouse preimplantation embryos. Mol. Reprod. Dev. 59, 277–284.

Bellier, S. (1997). Nuclear translocation and carboxyl-terminal domain phosphorylation of RNA polymerase II delineate the two phases of zygotic gene activation in mammalian embryos. EMBO J. 16, 6250–6262.

Bensaude, O. (2011). Inhibiting eukaryotic transcription. Which compound to choose? How to evaluate its activity?: Which compound to choose? How to evaluate its activity? Transcription 2, 103–108.

Boisvert, F.-M., van Koningsbruggen, S., Navascués, J. and Lamond, A. I. (2007). The multifunctional nucleolus. Nat. Rev. Mol. Cell Biol. 8, 574–585.

Bonev, B. and Cavalli, G. (2016). Organization and function of the 3D genome. Nat. Rev. Genet. 17, 661–678.

Bonnet-Garnier, A., Feuerstein, P., Chebrout, M., Fleurot, R., Jan, H.-U., Debey, P. and Beaujean, N. (2013). Genome organization and epigenetic marks in mouse germinal vesicle oocytes. Int. J. Dev. Biol. 56, 877–887.

Bonnet-Garnier, A., Kiêu, K., Aguirre-Lavin, T., Tar, K., Flores, P., Liu, Z., Peynot, N., Chebrout, M., Dinnyés, A., Duranthon, V., et al. (2018). Three-dimensional analysis of nuclear heterochromatin distribution during early development in the rabbit. Chromosoma 127, 387–403.

Bruno, P. M., Lu, M., Dennis, K. A., Inam, H., Moore, C. J., Sheehe, J., Elledge, S. J., Hemann, M. T. and Pritchard, J. R. (2020). The primary mechanism of cytotoxicity of the chemotherapeutic agent CX-5461 is topoisomerase II poisoning. Proc. Natl. Acad. Sci. 117, 4053–4060.

Bui, L. C., Evsikov, A. V., Khan, D. R., Archilla, C., Peynot, N., Hénaut, A., Le Bourhis, D., Vignon, X., Renard, J. P. and Duranthon, V. (2009). Retrotransposon expression as a defining event of genome reprograming in fertilized and cloned bovine embryos. REPRODUCTION 138, 289– 299.

Burns, K. H., Viveiros, M. M., Ren, Y., Wang, P., DeMayo, F. J., Frail, D. E., Eppig, J. J. and Matzuk, M. M. (2003). Roles of NPM2 in chromatin and nucleolar organization in oocytes and embryos. Science 300, 633–6.

Casanova, M., Pasternak, M., El Marjou, F., Le Baccon, P., Probst, A. V. and Almouzni, G. (2013). Heterochromatin Reorganization during Early Mouse Development Requires a Single-Stranded Noncoding Transcript. Cell Rep. 4, 1156–1167.

Cremer, T. and Cremer, C. (2001). Chromosome territories, nuclear architecture and gene regulation in mammalian cells. Nat Rev Genet 2, 292–301.

Drygin, D., Lin, A., Bliesath, J., Ho, C. B., O’Brien, S. E., Proffitt, C., Omori, M., Haddach, M., Schwaebe, M. K., Siddiqui-Jain, A., et al. (2011). Targeting RNA Polymerase I with an Oral Small Molecule CX-5461 Inhibits Ribosomal RNA Synthesis and Solid Tumor Growth. Cancer Res. 71, 1418–1430.

Fléchon, J.-E. and Kopecny, V. (1998). The nature of the ‘nucleolus precursor body’ in early preimplantation embryos: a review of fine-structure cytochemical, immunocytochemical and autoradiographic data related to nucleolar function. Zygote 6, 183–191.

Fulka, H. and Aoki, F. (2016). Nucleolus Precursor Bodies and Ribosome Biogenesis in Early Mammalian Embryos: Old Theories and New Discoveries1. Biol. Reprod. 94,.

Fulka, H. and Langerova, A. (2014). The maternal nucleolus plays a key role in centromere satellite maintenance during the oocyte to embryo transition. Development 141, 1694–1704.

Fulka, H. and Langerova, A. (2019). Nucleoli in embryos: a central structural platform for embryonic chromatin remodeling? Chromosome Res. 27, 129–140.

Gelali, E., Girelli, G., Matsumoto, M., Wernersson, E., Custodio, J., Mota, A., Schweitzer, M., Ferenc, K., Li, X., Mirzazadeh, R., et al. (2019). iFISH is a publically available resource enabling versatile DNA FISH to study genome architecture. Nat. Commun. 10, 1636.

Grummt, I. (2013). The nucleolus—guardian of cellular homeostasis and genome integrity. Chromosoma 122, 487–497.

Grummt, I. and Längst, G. (2013). Epigenetic control of RNA polymerase I transcription in mammalian cells. Biochim. Biophys. Acta BBA-Gene Regul. Mech. 1829, 393–404.

Guenatri, M., Bailly, D., Maison, C. and Almouzni, G. (2004). Mouse centric and pericentric satellite repeats form distinct functional heterochromatin. J. Cell Biol. 166, 493–505.

Guetg, C., Lienemann, P., Sirri, V., Grummt, I., Hernandez-Verdun, D., Hottiger, M. O., Fussenegger, M. and Santoro, R. (2010). The NoRC complex mediates the heterochromatin formation and stability of silent rRNA genes and centromeric repeats. EMBO J 29,.

Hamatani, T., Carter, M. G., Sharov, A. A. and Ko, M. S. H. (2004). Dynamics of global gene expression changes during mouse preimplantation development. Dev Cell 6,.

Hamatani, T., Ko, M. S., Yamada, M., Kuji, N., Mizusawa, Y., Shoji, M., Hada, T., Asada, H., Maruyama, T. and Yoshimura, Y. (2006). Global gene expression profiling of preimplantation embryos. Hum. Cell 19, 98–117.

Hamdane, N., Stefanovsky, V. Y., Tremblay, M. G., Németh, A., Paquet, E., Lessard, F., Sanij, E., Hannan, R. and Moss, T. (2014). Conditional Inactivation of Upstream Binding Factor Reveals Its Epigenetic Functions and the Existence of a Somatic Nucleolar Precursor Body. PLoS Genet. 10, e1004505.

Hamdane, N., Tremblay, M. G., Dillinger, S., Stefanovsky, V. Y., Németh, A. and Moss, T. (2016). Disruption of the UBF gene induces aberrant somatic nucleolar bodies and disrupts embryo nucleolar precursor bodies. Gene.

Henras, A. K., Plisson-Chastang, C., O’Donohue, M.-F., Chakraborty, A. and Gleizes, P.-E. (2015). An overview of pre-ribosomal RNA processing in eukaryotes. WIREs RNA 6, 225–242.

Herdman, C., Mars, J.-C., Stefanovsky, V. Y., Tremblay, M. G., Sabourin-Felix, M., Lindsay, H., Robinson, M. D. and Moss, T. (2017). A unique enhancer boundary complex on the mouse ribosomal RNA genes persists after loss of Rrn3 or UBF and the inactivation of RNA polymerase I transcription. PLoS Genet. 13, e1006899.

Inoue, A. and Aoki, F. (2010). Role of the nucleoplasmin 2 C-terminal domain in the formation of nucleolus-like bodies in mouse oocytes. FASEB J. Off. Publ. Fed. Am. Soc. Exp. Biol. 24, 485– 494.

Jachowicz, J. W., Santenard, A., Bender, A., Muller, J. and Torres-Padilla, M.-E. (2013). Heterochromatin establishment at pericentromeres depends on nuclear position. Genes Dev. 27, 2427–2432.

Jansz, N. and Torres-Padilla, M.-E. (2019). Genome activation and architecture in the early mammalian embryo. Curr. Opin. Genet. Dev. 55, 52–58.

Junera, H. R., Masson, C., Geraud, G. and Hernandez-Verdun, D. (1995). The three-dimensional organization of ribosomal genes and the architecture of the nucleoli vary with G1, S and G2 phases. J. Cell Sci. 108, 3427–3441.

Kent, T., Lapik, Y. R. and Pestov, D. G. (2008). The 5’ external transcribed spacer in mouse ribosomal RNA contains two cleavage sites. RNA 15, 14–20.

Koné, M. C., Fleurot, R., Chebrout, M., Debey, P., Beaujean, N. and Bonnet-Garnier, A. (2016). Three-Dimensional Distribution of UBF and Nopp140 in Relationship to Ribosomal DNA Transcription During Mouse Preimplantation Development1. Biol. Reprod. 94,.

Kyogoku, H., Fulka, J., Wakayama, T. and Miyano, T. (2014). De novo formation of nucleoli in developing mouse embryos originating from enucleolated zygotes. Development 141, 2255– 2259.

Lavrentyeva, E., Shishova, K., Kagarlitsky, G. and Zatsepina, O. (2015). Localisation of RNAs and proteins in nucleolar precursor bodies of early mouse embryos. Reprod. Fertil. Dev.

Le Bouteiller, M., Souilhol, C., Beck-Cormier, S., Stedman, A., Burlen-Defranoux, O., Vandormael-Pournin, S., Bernex, F., Cumano, A. and Cohen-Tannoudji, M. (2013). Notchless-dependent ribosome synthesis is required for the maintenance of adult hematopoietic stem cells. J. Exp. Med. 210, 2351–2369.

Lehnertz, B., Ueda, Y., Derijck, A. A., Braunschweig, U., Perez-Burgos, L., Kubicek, S., Chen, T., Li, E., Jenuwein, T. and Peters, A. H. (2003). Suv39h-mediated histone H3 lysine 9 methylation directs DNA methylation to major satellite repeats at pericentric heterochromatin. Curr. Biol. 13, 1192–1200.

Maiser, A., Dillinger, S., Längst, G., Schermelleh, L., Leonhardt, H. and Németh, A. (2020). Super-resolution in situ analysis of active ribosomal DNA chromatin organization in the nucleolus. Sci. Rep. 10, 7462.

Mangan, H., Gailín, M. Ó. and McStay, B. (2017). Integrating the genomic architecture of human nucleolar organizer regions with the biophysical properties of nucleoli. FEBS J. 284, 3977– 3985.

Mars, J.-C., Tremblay, M. G., Valere, M., Sibai, D. S., Sabourin-Felix, M., Lessard, F. and Moss, T. (2020). The chemotherapeutic agent CX-5461 irreversibly blocks RNA polymerase I initiation and promoter release to cause nucleolar disruption, DNA damage and cell inviability. NAR Cancer 2,.

Mayer, R., Brero, A., von Hase, J., Schroeder, T., Cremer, T. and Dietzel, S. (2005). Common themes and cell type specific variations of higher order chromatin arrangements in the mouse. BMC Cell Biol. 6, 44.

Meaburn, K. J. (2016). Spatial Genome Organization and Its Emerging Role as a Potential Diagnosis Tool. Front. Genet. 7,.

Miyanari, Y. and Torres-Padilla, M. E. (2012). Control of ground-state pluripotency by allelic regulation of Nanog. Nature 483, 470–3.

Moss, T., Mars, J.-C., Tremblay, M. G. and Sabourin-Felix, M. (2019). The chromatin landscape of the ribosomal RNA genes in mouse and human. Chromosome Res. 27, 31–40.

Mullineux, S.-T. and Lafontaine, D. L. J. (2012). Mapping the cleavage sites on mammalian pre-rRNAs: Where do we stand? Biochimie 94, 1521–1532.

Ogushi, S. and Saitou, M. (2010). The nucleolus in the mouse oocyte is required for the early step of both female and male pronucleus organization. J. Reprod. Dev. 56, 495–501.

Ogushi, S., Palmieri, C., Fulka, H., Saitou, M., Miyano, T. and Fulka, J. (2008). The Maternal Nucleolus Is Essential for Early Embryonic Development in Mammals. Science 319, 613.

Ogushi, S., Yamagata, K., Obuse, C., Furuta, K., Wakayama, T., Matzuk, M. M. and Saitou, M. (2017). Reconstitution of the oocyte nucleolus in mice through a single nucleolar protein, NPM2. J. Cell Sci. 130, 2416–2429.

Padeken, J. and Heun, P. (2014). Nucleolus and nuclear periphery: Velcro for heterochromatin. Curr. Opin. Cell Biol. 28, 54–60.

Pederson, T. (2011). The Nucleolus. Cold Spring Harb. Perspect. Biol. 3, a000638–a000638.

Perry, R. P. and Kelley, D. E. (1970). Inhibition of RNA synthesis by actinomycin D: characteristic dose-response of different RNA species. J. Cell. Physiol. 76, 127–139.

Potapova, T. A. and Gerton, J. L. (2019). Ribosomal DNA and the nucleolus in the context of genome organization. Chromosome Res. 27, 109–127.

Probst, A. V., Santos, F., Reik, W., Almouzni, G. and Dean, W. (2007). Structural differences in centromeric heterochromatin are spatially reconciled on fertilisation in the mouse zygote. Chromosoma 116,.

Probst, Aline. V., Okamoto, I., Casanova, M., El Marjou, F., Le Baccon, P. and Almouzni, G. (2010). A Strand-Specific Burst in Transcription of Pericentric Satellites Is Required for Chromocenter Formation and Early Mouse Development. Dev. Cell 19, 625–638.

Romanova, L. (2006). High Resolution Mapping of Ribosomal DNA in Early Mouse Embryos by Fluorescence In Situ Hybridization. Biol. Reprod. 74, 807–815.

Sanij, E., Hannan, K. M., Xuan, J., Yan, S., Ahern, J. E., Trigos, A. S., Brajanovski, N., Son, J., Chan, K. T., Kondrashova, O., et al. (2020). CX-5461 activates the DNA damage response and demonstrates therapeutic efficacy in high-grade serous ovarian cancer. Nat. Commun. 11, 2641.

Savić, N., Bär, D., Leone, S., Frommel, S. C., Weber, F. A., Vollenweider, E., Ferrari, E., Ziegler, U., Kaech, A., Shakhova, O., et al. (2014). lncRNA Maturation to Initiate Heterochromatin Formation in the Nucleolus Is Required for Exit from Pluripotency in ESCs. Cell Stem Cell 15, 720–734.

Schindelin, J., Arganda-Carreras, I., Frise, E., Kaynig, V., Longair, M., Pietzsch, T., Preibisch, S., Rueden, C., Saalfeld, S., Schmid, B., et al. (2012). Fiji: an open-source platform for biological-image analysis. Nat. Methods 9, 676–682.

Schindelin, J., Rueden, C. T., Hiner, M. C. and Eliceiri, K. W. (2015). The ImageJ ecosystem: An open platform for biomedical image analysis. Mol. Reprod. Dev. 82, 518–529.

Schneider, C. A., Rasband, W. S. and Eliceiri, K. W. (2012). NIH Image to ImageJ: 25 years of image analysis. Nat. Methods 9, 671–675.

Schöfer, C., Weipoltshammer, K., Almeder, M., Müller, M. and Wachtler, F. (1996). Redistribution of ribosomal DNA after blocking of transcription induced by actinomycin D. Chromosome Res. 4, 384–391.

Shishova, K. V. and others (2011). The fate of the nucleolus during mitosis: comparative analysis of localization of some forms of pre-rRNA by fluorescent in situ hybridization in NIH/3T3 mouse fibroblasts. Acta Naturae Англоязычная Версия 3,.

Shishova, K. V., Lavrentyeva, E. A., Dobrucki, J. W. and Zatsepina, O. V. (2015a). Nucleolus-like bodies of fully-grown mouse oocytes contain key nucleolar proteins but are impoverished for rRNA. Dev. Biol. 397, 267–281.

Shishova, K. V., Khodarovich, Y. M., Lavrentyeva, E. A. and Zatsepina, O. V. (2015b). High-resolution microscopy of active ribosomal genes and key members of the rRNA processing machinery inside nucleolus-like bodies of fully-grown mouse oocytes. Exp. Cell Res. 337, 208–218.

Solovei, I., Kreysing, M., Lanctôt, C., Kösem, S., Peichl, L., Cremer, T., Guck, J. and Joffe, B. (2009). Nuclear Architecture of Rod Photoreceptor Cells Adapts to Vision in Mammalian Evolution. Cell 137, 356–368.

Stalling, D., Westerhoff, M. and Hege, H.-C. Amira - a Highly Interactive System for Visual Data Analysis. 18.

Stefanovsky, V. Y., Pelletier, G., Bazett-Jones, D. P., Crane-Robinson, C. and Moss, T. (2001). DNA looping in the RNA polymerase I enhancesome is the result of non-cooperative in-phase bending by two UBF molecules. Nucleic Acids Res. 29, 3241–3247.

Szabo, Q., Donjon, A., Jerković, I., Papadopoulos, G. L., Cheutin, T., Bonev, B., Nora, E. P., Bruneau, B. G., Bantignies, F. and Cavalli, G. (2020). Regulation of single-cell genome organization into TADs and chromatin nanodomains. Nat. Genet. 52, 1151–1157.

van de Nobelen, S., Rosa-Garrido, M., Leers, J., Heath, H., Soochit, W., Joosen, L., Jonkers, I., Demmers, J., van der Reijden, M., Torrano, V., et al. (2010). CTCF regulates the local epigenetic state of ribosomal DNA repeats. Epigenetics Chromatin 3, 19.

van Sluis, M. and McStay, B. (2017). Nucleolar reorganization in response to rDNA damage. Curr. Opin. Cell Biol. 46, 81–86.

van Steensel, B. and Furlong, E. E. M. (2019). The role of transcription in shaping the spatial organization of the genome Nat. Rev. Mol. Cell Biol. 20, 327–337.

Wang, Q. T., Piotrowska, K., Ciemerych, M. A., Milenkovic, L., Scott, M. P., Davis, R. W. and Zernicka-Goetz, M. (2004). A Genome-Wide Study of Gene Activity Reveals Developmental Signaling Pathways in the Preimplantation Mouse Embryo. Dev. Cell 6, 133–144.

Zatsepina, O., Baly, C., Chebrout, M. and Debey, P. (2003). The Step-Wise Assembly of a Functional Nucleolus in Preimplantation Mouse Embryos Involves the Cajal (Coiled) Body. Dev. Biol. 253, 66–83.

Zentner, G. E., Balow, S. A. and Scacheri, P. C. (2014). Genomic Characterization of the Mouse Ribosomal DNA Locus. G3amp58 GenesGenomesGenetics 4, 243–254.

